# Adherence to Iron and Folic Acid Supplement and Associated Factors among Antenatal Care Attendant Mothers In Lay Armachiho Health Centers, Northwest, Ethiopia, 2017

**DOI:** 10.1101/493916

**Authors:** Gashaw Agegnehu, Azeb Atenafu, Henok Dagne, Baye Dagnew

## Abstract

**Background:** Iron deficiency is the leading nutrient deficiency in the world affecting the lives of more than 2 billion people, accounting to over 30% of the world’s population. Pregnant women are particularly at high risk of iron and folic acid deficiency.

**Objective:** The aim of this study was to assess Adherence to Iron and folic acid supplement during pregnancy and its associated factors among pregnant women attending antenatal care.

**Methods:** Institution based cross-sectional study was employed from February 2016 to March 2017. Systematic random sampling technique was used to select the study participants. Data was collected using a structured and pretested interviewer-administered questionnaire. Bivariable and multivariable logistic regression analysis were used to identify associated factors with Adherence to prenatal iron and folic acid supplement among pregnant women. An adjusted odds ratio with a 95% confidence interval was computed to determine the level of significance. Those variables with a p-value less than 0.05 had been considered as significant.

**Result:** Adherence to Iron and folic acid was 28.7% with 95% C.I. (24.3, 33.6%). Educational status of mothers(AOR= 9.27 (95%CI: 2.47, 34.71), Educational status of husband (AOR= 0.31(95% CI: 0.11,0.88), Mothers who had a family size of four(AOR=3.70(1.08,12.76), Mothers who had family size of five and above (AOR= 4.88(95% CI: 1.20, 19.85),Mothers who had 2500-3500 birr household average monthly income (AOR= 0.46(95% CI: 0.24,0.89), Mothers who had registered at 17-24weeks with (AOR=0.40(95% CI: 0.22-0.74), registered at 25-28weeks (AOR=0.20(95% CI 0.10, 0.41), Mothers who had collected their iron and folic acid started at first visit at first month of pregnancy and duration of iron and folic acid is taken (AOR= 2.42(95% CI:1.05, 5.58) had significant association with iron and folic acid adherence.

**Conclusion and recommendation:** Adherence of Iron and folic acid was relatively low. Maternal and husband education status, family size, registration time, economic status and first visit in the first month with duration of iron and folic acid taken were factors significantly associated with adherence to iron and folic acid supplement. Educating pregnant mothers, improving economic status, early ANC registration can improve adherence to iron and folic acid supplement.

## 1. Background

Iron deficiency is the leading single nutrient deficiency in the world affecting the lives of more than 2 billion people, accounting to over 30% of the world’s population particularly in developing countries [1]. Pregnant women are particularly at high risk of iron and folic acid deficiency due to increased nutrient requirement during pregnancy [2, 3]. In Ethiopia, 17% of reproductive age women are estimated to be anaemic and particularly 22% of pregnant women are anaemic from which 18% are found in rural and 11% are in urban residents [4, 5]. According to WHO and Ethiopia’s national guidelines for control and prevention of micronutrient deficiencies, all pregnant women should receive and consume a standard dose of 60mg iron and 400 μg folic acid daily for 6 months starting from first month of pregnancy or at the time of their first antenatal visit and three months of postnatal period [6–9].

Iron is a trace mineral that is vital for fetal growth and development, it plays a key role as a cofactor for enzymes involved in the oxidation-reduction reaction. Iron is found in red blood cells to carry oxygen needed throughout the body. Iron is also essential for normal neuronal development. Folic acid is another important micronutrient used in the synthesis of neurotransmitters and particularly during early pregnancy. It has an essential role in synthesizing DNA during organogenesis [10].

Iron and folic acid supplementation are required to balance increased physiological demand - during puberty, pregnancy, and lactation [6, 7]. This significantly increased demand for iron and folic acid is required for the development of fetus and placenta as well as in supporting maternal blood volume. Pregnant women require a much higher amount of iron and folic acid that is met by most diets and therefore to fulfil this they routinely receive iron and folic acid supplements. Pregnant women, postpartum women and children aged 6-24 months are usually the most affected groups [6, 9]. Iron and folic acid deficiency are higher in developing countries which can be contributed by malnutrition, parasitic and bacterial infections.

The high physiological requirement for iron and folic acid in pregnancy is difficult to meet with the usual diet. Therefore pregnant women should routinely receive iron and folic acid supplements. Adherence to medication regimen is generally defined as the extent to which patients take medications as prescribed by their health care providers [11]. Adherence rates for individual patients are usually reported as the percentage of the prescribed doses of the medication actually taken by the patient over a specified period [12].

One in every three women had anaemia while one in every two had iron and folic acid deficiency, indicating that both folic acid and iron deficiencies constitute the major micronutrient deficiencies in Ethiopian women [2].

Iron and folic acid deficiency is a serious public health issue due to its high potential negative consequences. It can lead to several adverse outcomes including low birth weight, preterm delivery, stillbirth, and maternal and neonatal mortality. Infants are among the vulnerable groups of iron/folic acid deficiency. Since there is a link between maternal and neonatal iron status, interventions on infant alone are insufficient to decrease infant iron and folic acid status. Oral iron and folic acid supplementation is a feasible and cost-effective strategy in prevention and control of iron and folic acid deficiency anaemia [13].

The risk of death decreases by 24% and 1.8 million deaths in children aged 28 days to 10 years will be avoided for each increase in 1g/dl haemoglobin. Simple IFAS strategies are feasible ways to attain such increments in haemoglobin [13]. Early neonatal death was reduced by 57% in Nepal and 45% in Pakistan when more than 90 tablets are taken and started at or before fifth months of pregnancy. Similarly, the risk was decreased by more than half in those study participants who took IFAS during pregnancy [14]. Currently, iron and folic acid supplementation is the main strategy for anaemia control and prevention in Ethiopia [5, 8]. While many developing countries including Ethiopia are now implementing IFAS through antenatal care programs, only a few countries have reported significant improvement in IFAS and anaemia control and prevention [5, 15, 16].

This low achievement may be related to poor access to and utilization of antenatal care services, inadequate supply of IFA tablets, poor counselling, lack of knowledge on anaemia, and certain belief [5]. However, many studies suggested poor maternal adherence to the regimen is the main reason for the ineffectiveness of the strategy [11, 15]. There is little feedback about the effectiveness of iron and folic acid supplement countrywide and there is concern that women would have poor adherence due to perceived side effects, particularly of iron supplements.

Therefore, the aim of this study is to assess the magnitude and associated factors of the adherence of iron/folic acid supplementationation among pregnant women in Layaremachiho district health centres, Northwest, Ethiopia.

## Methods

### Study design and settings

An institution based cross-sectional study design was employed from February to March 2017 in Lay Armachiho district Health Centers. Lay Armachiho district is one of 22 districts in North Gondar administrative zone and is located at a distance of 769 ***km*** from Addis Ababa, 202***km*** from Bahirdar and 22 ***km*** from Gondar Administrative city. It encompasses about 129,272 ***km^2^*** area of land and is characterized by diverse climate and topography with a marked difference in the climate conditions. Most of the area is on the highland plateau and is characterized by rugged mountains, hills and plateaus. Hence, the district has varied landscapes composed of steep fault escarpments and nearly mountainous, and erodible landforms but draught not occurred and no shortage of rain in a rainy season. It is located in North West of Gondar, Ethiopia and bordered by by Wogera, Sanja, Chillga, Dembia and North Gondar Administrative city. According to figures from the Central Statistical Agency, 2007, the district has an estimated population of 140, 417, of whom 70, 911 are females, Estimated pregnant women are 5,616. There are 6 health centers, 26 health posts, 6 junior private clinics, 1 Medium private clinic and 2 private Drug venders providing health services in the district [17].

### Sample size determination

The sample size was calculated by using single population proportion formula with the following assumptions: p (Proportion of Adherence to Iron and folic acid supplement =55.5 % [18], 95% confidence interval, and 5% margin of error (d).

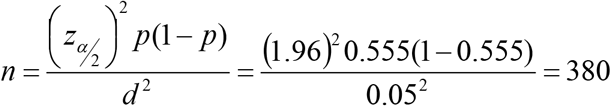

Considering, a 10% non-response rate, the final sample size was 418.

### Sampling procedures

There were a total of 838 pregnant women who fulfilled eligibility criteria during the study period in Lay Armachiho district, 2017. Using baseline information proportional number of study subjects was selected from each Health Center. Systematic random sampling technique was used to include a sample of 418 study participants.

### Data collection procedures

A structured questionnaire was used to collect data. The questionnaire was pre-tested over 30 women out of the study area in an institution with similar characteristics prior to the actual data collection. Six BSc Nurses (worked as data collectors) and two BSc Public Health professionals (worked as supervisors) were involved in the data collection process. Training was given for the data collectors and supervisors. The overall interview process was supervised by supervisors. The collected data were checked by the data collectors immediately after finalizing the questionnaire before they left the health centres. Supervisors daily checked the completeness and consistency of information collected.

### Measurement of study variables

Adherence to iron and folic acid supplement is the outcome variable in this study. Pregnant mothers are said to be “adherent” to Iron and folic acid supplement if they took 65% or more of the supplement, equivalent to taking the supplement at least 4 days a week during 3 months period using recording, reporting and checking their cards [19]. Early registration to ANC clinic was measured as those pregnant women who visit the ANC clinic before 16 weeks of gestation [20]. Knowledge to anaemia was measured by asking knowledge questions. Mothers who score median and more questions on anaemia are said to be knowledgeable [21]. Knowledge about iron and folic acid supplement was measured by asking several questions about IFAS. Those who score median and above on questions prepared to assess comprehensive knowledge of IFAS of the respondents [20].

### Data management and statistical analysis

Data were entered using EPI-INFO version 3.5.11 and exported into SPSS version 20 for analysis. For most variables, data were presented by frequencies and percentages. Univariable binary logistic regression analysis was used to explore candidate variables for the multivariable logistic regression analysis and variables with p-value less than 0.2 by the univariable analysis were then analyzed by multivariable logistic regression for controlling the possible effect of confounders and finally variables which had significant association with knowledge and practice were identified on the basis of AOR with 95% CI and p < 0.05. Hosmer and Lemeshow goodness of fit test was used to check model fitness. VIF was also used to test interactions between variables.

## Results

### Socio-demographic information

A total of 418 study participants participated in this study with 100 % response rate. The mean age of respondents was 27.2 (SD± 6.3) years. Majority of respondents 312 (74.6 %) were ruler dwellers. Four hundred fifteen (99.3%) of the study participants were followers of Orthodox Christianity. The majority (83%) of the study subjects were housewives (Table 1).

**Table 1:** Socio-Demographic factors of respondents, Lay Armachiho district Health Centers, Northwest Ethiopia, 2017.

| Variable |  | Frequency | Percent |
| --- | --- | --- | --- |
| Age | 17- 25 | 211 | 50.5 |
|  | 26- 35 | 155 | 37.1 |
|  | 36- 45 | 52 | 12.4 |
| Residence | Rural | 312 | 74.6 |
|  | Urban | 106 | 24.4 |
| Religion | Orthodox | 415 | 99.3 |
|  | Muslim | 3 | 0.3 |
| Ethnicity | Kemant | 380 | 90.9 |
|  | Amhara | 38 | 9.1 |
| Educational status | Cannot read and write | 188 | 45 |
|  | Completed primary and above | 230 | 55 |
| Educational status of husband | Cannot read and write | 118 | 28.2 |
|  | Completed primary and above | 300 | 71.8 |
| Occupation of mother | House wife | 347 | 83 |
|  | Government employee | 46 | 11 |
|  | Merchant | 25 | 6 |
| Occupation of husband | Government employee | 66 | 15.8 |
|  | Merchant | 56 | 13.4 |
|  | Farmer | 296 | 70.8 |
| Family size | Two | 101 | 24.2 |
|  | Three | 77 | 18.4 |
|  | Four | 64 | 15.3 |
|  | Five and above | 176 | 42.1 |
| Monthly income | 500- 2500 birr | 165 | 39.5 |
|  | 2501- 3500 birr | 161 | 38.5 |
|  | >3500 birr | 92 | 22 |
| Knowledge about iron foliate and anemia | Knowledgeable | 57 | 13.6 |
|  | Non-knowledgeable | 361 | 86.4 |

### Pregnancy, Obstetric, and health-related factors of respondents

Majority of respondents (72.2%) had less than three times ANC visit. Among the respondents, 124(29.7%) had started ANC during the first trimester (before 16 weeks of gestation)(Table 2).

### Respondents knowledge of anaemia and benefit of Iron and folic acid supplement

Less than one-fifth of respondents 57 (13.6%) were knowledgeable about anaemia, iron and folic acid supplementation, while 361(86.4%) of respondents were not knowledgeable about anaemia and iron and folic acid supplementation (Table 3).

### Health service-related factors of respondents, Lay Armachiho district health centres, Northwest Ethiopia, 2017

Only 4 study participants have used iron and folic acid for greater than three months (90 tablets). Among the respondents, 80% were provided health education about iron and folic acid supplements (Table 4).

### The proportion of adherence to Iron and folic acid supplementation

From a total of 418 participants included in this study, only120 (28.7%) participants adhered to iron-folic acid supplementation.

### Factors associated with adherence to iron and folic acid supplementation

Univariable binary logistic regression was used to choose variables for the final model on the basis of p-value less than 0.2. Place of residence, educational status of, educational status of husband, number of pregnancy, number of delivery, knowledge on anaemia, occupation of mother, occupation of husband, family size, early registration, average household monthly income, disease confirmed by physician like gastritis, treatment of disease and first visit at first month, were variables selected for the final model. VIF was calculated considering one independent variable as the dependent variable turn by turn to test interactions between variables. The test result shows that VIF for all variables was below 3, the threshold for co-linearity diagnostics. This showed that there is no multicolinearity effect between independent variables. Table 5 shows variables associated with adherence with **IFAS**. Adherence to IFAS was significantly associated with the educational status of participants. The probability of adherence to IFAS was 9.27 times higher among participants whose educational status was primary education and above [AOR= 9.27, 95%CI= (2.47, 34.71)] as compared to study participants who cannot read and write. About 69% of study participants whose husbands have an educational status above primary education were less likely adhered to IFAS as compared to those study subjects whose husbands cannot read and write [AOR= 0.31, 95% CI = (0.11, 0.89)]. The probability of IFAS adherence was 3.70 times higher among participants who have a family size of four as compared to those having only two family members[AOR=3.70, 95% C.I = (1.08,12.76)]. Adherence of IFAS among study participants with five and above family members was 4.88 times higher than those only with 2 family members [AOR=4.88, 95% C.I = (1.20, 19.85]. As revealed by this study, Adherence to IFAS was associated with average monthly household income. The study participants whose average monthly income is 2501 to 3500 ETB were less likely adhered to IFAS compared to those who earned 500 to 2500 ETB [AOR= 0.46, 95% CI = (0.24,0.90)]. Participants who had registered 17-24 weeks of gestation [AOR=0.40, 95% CI (0.22-0.74)] and participants who had registered at 25-28weeks [AOR=0.20, 95% CI (0.10, 0.41)] were less likely to adhere to IFAS. The probability of Adherence to IFAS was 2.42 times higher among study participants collecting their pill at first visit at first month of pregnancy with duration of pill used [AOR= 2.42, 95% CI (1.05, 5.58)].

## DISCUSSION

Adherence to iron and folic acid supplement found in this study was 28.7% with 95% C.I [24.3-33.6]. This is lower than studies conducted in Cambodia [14], India and Indonesia [13]. It is also lower than studies done among pregnant women in Eritrean refugee camps, Northern Ethiopia [22], among pregnant women attending ANC at University of Gondar hospital [23], and Governmental Health Centers in Akaki Kality Sub City [24]. But the adherence in the current study is higher than results from studies conducted in Kenya[25], Goba, Ethiopia[18], **a** study done in 8 rural districts of Ethiopia [21], and a study conducted in Mecha district in Amhara region [26]. This difference could be due to the differences in socioeconomic status of the study populations, present study based on rural population and the time of the study.

This study depicted that educational status of the mother was significantly associated with adherence to IFAS. The adherence was higher among mothers who have completed primary education and above. This finding is in line with the findings of other studies in Ethiopia [26] and India [27]. Education is more likely to enhance female awareness of micronutrient deficiency and ways to overcome these deficiencies. Overall, educated women have greater ability to stick to health care inputs such as IFA which offer better care for both the infant and the mother.

A husband who completed primary education and above were less likely promote adherence of iron-folic acid supplementation. This result is inconsistent with a study conducted in Indonesia both mother and fathers had promoted and enhanced adherence status of pregnant women equally [28]. Household average monthly income was associated with iron and folic acid supplementation. This was consistent with a study conducted in Kenya[25]. Family size was significantly associated with adherence of IFAS. This might be because those participants who had multiparty would have medical advice, health education, counselling and access to more information about the benefits of iron and folic acid during pregnancy and early pregnancy than nulliparous participants.

The current study has revealed that those participants with lower family income are less likely to adhere with iron and folic acid supplementation compared to those with higher family income. Early time of registration was significantly associated with IFAS adherence. This finding is consistent with a study done in Kenya[25].

## Conclusion

The adherence of IFAS among antenatal care attendant women in Lay Armachiho district health centres was found to be a low and significant proportion of women do not adhere to IFAS. Educational status of participants, educational status of husband, family size, household average monthly income, registered time and tablet collection time were identified as factors associated with adherence to IFAS. Educating pregnant mothers, improving economic status and early ANC registration can improve adherence to IFAS.

## List of abbreviations

ANC: Antenatal care
AOR: Adjusted Odds Ratio
CI: Confidence Interval
COR: Crude Odds Ratio
IFAS: Iron-folic acid Supplementation
Km: Kilometer
Km^2^: Square kilometers
mg: milligram
SPSS: Statistical Package for Social Sciences
VIF: Variance inflation factor
WHO: World Health Organization
μg: microgram

## Declaration

### Ethics approval and consent to participate

Ethical clearance was obtained from the Institutional Review Board of the University of Gondar and an official letter was submitted to the health centres. There were no risks due to participation in this research project. The collected data were used for this research purpose only and kept with complete confidentiality. Verbal informed consent was obtained from the study participants Researchers provided health education for the study subjects about the importance of IFAS.

### Consent for publication

This manuscript does not contain any individual person’s data.

### Availability of data and material

Data will be made available upon request the primary author.

### Authors’ contribution

All the authors actively participated during the conception of the research issue, development of a research proposal, analysis and interpretation and writing various parts of the research report **GA** designed the protocol; participate in data collection; entered data into Epi-Info epidemiological software; analyzed the data and supervised the overall research process. **AA** designed the protocol and supervised the overall research process. **HD** designed the protocol, supervised the overall research process and prepared the manuscript. **BD** undertakes data analysis, commented on the manuscript and revised accordingly. All the authors read and approved the final manuscript.

## Acknowledgement

The authors are pleased to acknowledge data collectors, field supervisors, study participants, University of Gondar and managers and staff members of Lay Armachiho District Health Centers for their unreserved contributions to the success of this study.

